# *Oligella otitidis* sp. nov., isolated from middle ear discharge of children with chronic suppurative otitis media

**DOI:** 10.64898/2026.06.29.735399

**Authors:** Jemima Beissbarth, Brianna Atto, Pappu K Mandal, Alexander Cleanthous, Bailey Harrison, Nicholas Gill, Heidi C Smith-Vaughan, Mariana Kleinecke, Vanessa Rigas, Amanda J Leach, Peter S Morris, Robyn L Marsh

## Abstract

*Oligella otitidis* MSHR-50489EDL strain (ATCC: TSD-462-; DSMZ: DSM 118617) is a new species of the genus *Oligella* that was isolated from a middle ear discharge swab from a child with chronic suppurative otitis media (CSOM). This Gram-negative coccobacillus produces small, circular, smooth, whitish-opaque and occasionally mucoid colonies. It grows in aerobic conditions at a temperature range from 25-42°C. Phylogenetic analysis demonstrates a relationship to other species of the genera *Oligella* and average nucleotide identity and digital DNA–DNA hybridization values indicate a distinct species in comparison to other *Oligella* species. Thus far, the majority of isolates exhibit resistance to ciprofloxacin, the first line treatment for CSOM.

## Introduction

We report an addition to the genus *Oligella*. Previously two *Oligella* species have been described in the LPSN database: *O. urethralis* and *O. urealytica*^1^. These species are usually found in the human genitourinary tract. Occasionally, they have been isolated from other sites when causing serious infections among immunocompromised hosts^2^. Currently, there is limited understanding about *Oligella* and only a small number of published genomes^3^.

Bacteriome studies from Australia and the Philippines have detected *Oligella* reads in short-read 16S rRNA amplicon sequence data from ear discharge samples^4,5^; however, strains were not cultured. We previously cultured *Oligella urethralis* from the ear discharge of a child with chronic suppurative otitis media (CSOM)^6^, a severe form of middle ear infection. Here, we report isolation of a novel *Oligella* lineage from CSOM ear discharge that we propose be named *Oligella otitidis* sp. nov.

## Methods

### Isolation and growth conditions

Isolates (n=30) of a novel *Oligella* species (*Oligella otitidis* sp. nov.) were recovered from CSOM middle ear discharge swabs collected from children residing in the Northern Territory, Australia. Children were enrolled in a randomised controlled trial comparing CSOM treatments^7^, with informed consent provided for ear discharge sampling and microbiological testing. The study was approved by Human Research Ethics Committee of Northern Territory Health and Menzies School of Health Research (Approval #2014-2170). All swabs were stored in skim-milk tryptone glucose glycerol medium (STGGB) at -80°C prior to being cultured. Strains were isolated by inoculating STGGB medium onto MacConkey without salt agar (Thermo Scientific; pH 7.1) and incubating at 37°C in 5% CO_2_ for 48 hours. Isolates were then passaged onto horse blood Colombia agar (Thermo Scientific; pH 7.3) and grown in aerobic conditions at 37°C. Strains were not identified by MALDI-TOF mass spectrometry (MicroFlex spectrometer, Bruker, USA) but results were suggestive of *O. urethralis* with a MALDI-TOF quality confidence score range of 1.6-1.86.

### Phenotypic characterisation

To assess thermotolerance and aerotolerance of *O. otitidis* (n=30) isolates, horse blood agar (Thermo Scientific, Oxoid) plates were inoculated and incubated for 48 hours under aerobic conditions (4°C, 35°C, 37°C, 42°C or 60°C) or varying atmospheric conditions (aerobic, 5% CO_2,_ anaerobic [<1% O_2_ AnaeroGen, Thermo Scientific, Oxoid] or microaerophilic [8-9% O_2_/7-8% CO_2_; CampyGen, Thermo Scientific, Oxoid]) alongside control organisms *Pseudomonas aeruginosa* ATCC 15692 and *Campylobacter jejuni* ATCC 33560. Growth curves for isolates were generated to assess *O. otitidis* growth in liquid media (Mueller–Hinton broth, Heart Infusion broth and Tryptone Soya broth; Thermo Scientific), halotolerance, pH tolerance and effect of commonly used media supplements (1% tryptone [Thermo Scientific, Oxoid], 5% lysed horse blood [Thermo Scientific] and/or 20 mg/mL NAD [Merck]). Colonies of *O. otitidis* from 48-hour growth on horse blood agar were suspended to an OD_600_ = ∼0.1 (∼3 × 10^8^ CFU/mL) in sterile PBS and added to inner wells of a 96-well microtiter plate (in triplicate) to a total volume of 200 μL and ∼3-7 × 10^5^ CFU/mL. Plates were incubated at 37°C in 5% CO_2_ with 200 RPM orbital shaking in a Variskan LUX plate reader (Thermo Scientific) with OD_600_ measured every 30 minutes for 24-48 hours (or until end of log phase). Exit cultures were performed to detect contaminants by removing 5 μL from each well and spotting onto chocolate agar (Thermo Scientific, Oxoid). Biochemical profiling was determined using a VITEK2 NH card, as per the manufacturer’s instructions (BioMerieux).

### Antibiotic susceptibility

Due to the lack of *Oligella*-specific antibiotic susceptibility guidelines, susceptibility was assessed based on the CDS method using SensiTest agar (ThermoFisher Scientific) with an annular radius threshold of >6 mm arbitrarily applied. The MIC was determined by broth micro-dilution (EUCAST v7.0, Jan 2022) using cation adjusted Mueller Hinton broth.

### Genome sequencing

DNA was extracted from pure colonies (¼-of a 10μL sterile loop) suspended in 360 μL of enzymatic lysis buffer (20 mM Tris.Cl pH 8.0, 2mM EDTA, 1.2% Triton, 20 mg mL^-1^ lysozyme) using a QIAamp DNA Mini Kit bacterial protocol without modification^8^.

The type strain (*O. otitidis* MSHR-50489EDL) was sequenced using Illumina Novaseq 6000 (150-bp paired-end) by a commercial provider (AGRF, Australia) using Illumina DNA Prep PCR-Free library workflow (v1.5 chemistry, XP-lane splitter kit). Illumina sequencing generated 8,613,450 reads (473X coverage) which were filtered and trimmed using Trimmomatic v0.36^9^ (Settings - Leading: 20; Trailing: 20; Sliding window: 4:20; Minimum length: 30).

Long-read sequencing of all *O. otitidis* (n=30) isolates was performed using Oxford Nanopore MinION Mk1C using a FLO-MIN106(R 9.4) flowcell and rapid barcoding kit (SQK-RB114.96) for library preparation. Data acquisition was done using MinKNOW v22.8.9 and basecalling/demultiplexing was performed using Guppy v6.2.7 (Oxford Nanopore) in high-accuracy mode (pass = Q>9). A total of 2.37 Gb passed reads were generated with an average 110X coverage. Raw nanopore reads were processed using the EPI2ME wf-bacterial-genomes workflow (v2.0.1). Reads were concatenated and filtered using fastcat v0.22.1 (minimum read length 1000 bp, exclusion of low-complexity sequences [fastlint threshold = 20]). *De novo* assemblies were generated using Flye v2.9.6 using standard bacterial assembly mode (genome size ∼2-3 Mb, i=1 and single-genome bacterial assembly mode) and polished using Medaka v1.12.1, generating singular contig circular genomes 2.55-2.76 Mbp in length (N50 = 2.55-2.76 Mbp).

*De novo* hybrid assembly of MinION long reads and Illumina-filtered reads was done using Unicycler v0.5.0^10^. The origin of the genome was determined using Ori-Finder 2022^11^ and the genome was opened at *OriC* using Artemis v18.2.0^12^. *O. otitidis* MSHR-50489EDL (GCA_ CP158480) was deposited in ATCC (ATCC: TSD-462) and DSMZ (DSM 118617).

A total of 27 publicly available whole genome assemblies for *O. ureolytica* and *O. urethralis* were available from NCBI (accessed 02/03/2026). Genomes were assessed based on assembly size, contiguity, completeness, contamination, GC content and ANI to species type strains (*O. urethralis* NCTC 12964, GCA_900454345; *O. ureolytica* NCTC 11997, GCA_900454285). NCBI genomes were retained in downstream comparative analyses if they had genome sizes of 2.0–2.8 Mb, completeness ≥90%, contamination <10%, GC content 46% ± 0.7 and ANI ≥95-99% compared to the corresponding type strain (with <95% indicating incorrect species assignment and >99% indicating strain duplication). For *O. ureolytica*, slightly relaxed contamination thresholds were applied because otherwise high-quality assemblies showed uniformly elevated contamination estimates despite near-complete assembly status, low contig counts, consistent GC content and ANI values concordant with species assignment. While a more stringent approach to genome quality would be ideal (N50 >100 kb, contig number <50 and completeness of >95%), this would rule out most genomes, reflecting a poor *O. urealytica* genome representation in NCBI. A total of 17 genomes (3 *O. ureolytica* and 14 *O. urethralis*) were deemed acceptable for pangenome and phylogenetic analyses.

### Genome annotation

Annotation of local and NCBI genomes was performed using Prokka (v1.15.6) with default bacterial annotation settings. Annotated features were screened for common putative determinants of ciprofloxacin resistance in Gram-negative bacteria. Specifically, chromosomal quinolone target genes *gyrA* and *gyrB* (DNA gyrase subunits), and *parC* and *parE* (topoisomerase IV subunits; or synonymous *grlA* and *grlB*) were extracted for further analysis. Protein sequences corresponding to quinolone resistance-determining regions (QRDRs) were extracted from annotated genomes and aligned using MAFFT v7.526. QRDR boundaries were defined based on established coordinates in *Escherichia coli* K-12 substr. MG1655 (GCA_000005845.2) and subsequently refined through cross-species alignment to account for sequence divergence within the genus *Oligella*. Amino acid substitutions were identified relative to reference sequences and interpreted in the context of previously reported resistance-associated mutations in Gram-negative bacteria. While additional quinolone resistance mechanisms have been described, including pentapeptide repeat proteins, quinolone-modifying enzymes, and efflux-associated genes and regulators, the presence of these determinants alone is not reliably predictive of phenotypic resistance, as their contribution is often dependent on gene expression and regulatory context. Accordingly, analysis was restricted to QRDR mutations, which have well-established associations with quinolone MICs across Gram-negative bacteria.

### Comparative genomics

A core genome alignment was generated using Roary v3.13.0, applying a minimum BLASTp identity threshold of 85%. A maximum-likelihood phylogenetic tree was constructed using MEGA v12.0.11, applying the nucleotide substitution model selected based on model testing (General Time Reversible mode) and branch support was assessed using bootstrap analysis with 1000 replicates.

The degree of genomic similarity between *O. otitidis* sp. nov. and known *Oligella* spp. was determined by comparing percent average nucleotide identity (ANI), digital DNA-DNA hybridization (dDDH) value and average amino acid identity (AAI) with complete genomes of two *O. urethralis* ear strains (type strain NCTC 12964 [AKA DSM 7531/ATCC17960] GCA_ 900454345, and NCTC 11008 GCA_ 900454325) and two *O. ureolytica* urine isolates (type strain NCTC11997 [AKA DSM 18253/ATCC 43534] GCA_900454285, and NCTC 11998 GCA_900454315) using FastANI v1.34 (species boundary: ≥95–96%), GGDC v3.0^13,14^ (species boundary: ≥70%) and CompareM v0.1.2 (species boundary: ≥95–96%), respectively.

Pangenome analysis was performed to compare the genome of *O. otitidis* and NCBI whole genomes of other *Oligella* spp. using Roary v3.13.0 with a BLASTP-based ortholog clustering strategy and core gene alignment using MAFFT v7.526. Orthologous gene clusters were defined based on amino acid sequence similarity, and a gene presence–absence matrix was generated for all analysed 30 *O. otitidis* genomes and 17 NCBI genomes (≥95% threshold). Genes uniquely associated with the new taxa were defined as those present in *O. otitidis* sp. nov. genomes (n=30) and absent from *O. urethralis* and *O. ureolytica* NCBI genomes (n= 17).

### Phylogenetic analysis

Phylogeny was inferred using the Maximum Likelihood method and General Time Reversible model^15^ of nucleotide substitutions and the tree with the highest log-likelihood (-162,655.47) is shown. The tree is drawn to scale with branch lengths (shown below the branches) computed using the Maximum Likelihood method^15^ and measured in the number of substitutions per site. The percentage of replicate trees in which the associated taxa clustered together (1,000 replicates) is shown next to the branches^16^. The initial tree for the heuristic search was selected by choosing the tree with the superior log-likelihood between a Neighbor-Joining (NJ) tree^17^ and a Maximum Parsimony (MP) tree. The NJ tree was generated using a matrix of pairwise distances computed using the General Time Reversible model^15^. The MP tree had the shortest length among 10 MP tree searches, each performed with a randomly generated starting tree. The evolutionary rate differences among sites were modelled using a discrete Gamma distribution across 5 categories (+G, parameter = 0.4302), with 46.62% of sites deemed evolutionarily invariant (+I). The analytical procedure encompassed 47 nucleotide sequences. The complete deletion option was applied to eliminate positions containing gaps and missing data resulting in a final data set comprising 49,440 positions. Evolutionary analyses were conducted in MEGA12^18^ using up to 8 parallel computing threads.

## Results and Discussion

### Phenotypic characteristics

*Oligella otitidis* sp. nov is a Gram-negative, non-spore-forming cocco-bacillus that is oxidase- and catalase-positive, consistent with other members of the genus. Growth occurs on routine laboratory media, including horse blood and MacConkey agar, at 25–42 °C. Pinpoint colonies are observed after 24 hours incubation. After 48 hours incubation, the colonies appear small, circular, smooth, whitish-opaque and occasionally mucoid. Growth occurs aerobically (with or without 5% CO_2_) and under microaerophilic conditions, with little to no growth observed anaerobically. Growth is observed at pH 6–10 (optimum pH 8–9; no growth at pH 5), and at NaCl concentrations of 0.5–6.5% (optimum 4–5%) indicating moderate halotolerance. Growth was not recovered in tryptic soy broth supplemented with 1% tryptone, 5% lysed horse blood and/or 20 mg/mL NAD – suggesting a lack of dependence on these factors for growth.

To date, reports of phenotypic and biochemical characterisation of the *Oligella* genus are limited, with many characteristics either variably reported or not assessed across studies^3^, particularly for commercial identification systems. In this study, all *O. otitidis* sp. nov. isolates and *O. ureolytica* ATCC 43534 were identified as *O. urethralis* (probability 95-99%) by Vitek 2 NH cards, suggesting common biochemical characteristics across the genus (and limited utility of Vitek 2 NH cards for species-level identification). *O. otitidis* sp. nov. exhibited a strictly asaccharolytic metabolism, with all isolates negative for carbohydrate utilisation (including D-glucose, D-mannose, D-maltose, sucrose, D-xylose, glycogen and N-acetyl-D-glucosamine) which is consistent with other members of the genus^3^. Enzymatic activities indicate reliance on amino-acid and peptide metabolism, with positive reactions for multiple arylamidases and γ-glutamyl transferase. This is supported by the strong growth observed in heart infusion broth and restricted growth in tryptic soy broth, suggesting strong dependence on peptide-rich media and a preference for complex peptones or tissue digests. As with *O. urethralis*^*3*^, *O. otitidis* sp. nov. can be differentiated from *O. ureolytica* by the absence of urease activity and non-motility. Phenotypic characteristics and biochemical reactions of *O. otitidis* sp. nov., together with comparisons to the limited available data for other members of the genus, are summarised in Table 1. MALDI-TOF differentiated *Oligella* species when reference spectra for *O. otitidis* were made available in the library.

**Table 1:** Phenotypic/biochemical profile of 30 *Oligella otitidis* sp. nov. compared to reported *O. urethralis* (n= 118) and *O. ureolytica* (n=13) phenotypes^3^.

| Phenotype | *O. otitidis* sp. nov.<br>(%) | *O. urethralis* | *O. ureolytica* |
| --- | --- | --- | --- |
| Cell morphology | Gram-negative<br>coccobacilli | Gram-negative<br>coccobacilli | Gram-negative<br>coccobacilli |
| Motility | – (100) | – | + |
| Oxidase | + | + | + |
| Catalase | + | + | + |
| Growth on MacConkey | + | +/- | + |
| Beta-haemolysis | – (100) | – | – |
| Growth at 25 °C | + | v | NR |
| Growth at 35-37 °C | + | + | + |
| Growth at 42 °C | + | +/- | - |
| NaCl tolerance (4.5%) | + | +/- | + |
| Oxygen requirement | Aerobic (100) | Aerobic | Aerobic |
| pH growth range | 6-10 (100) | NR | NR |
| Vitek NH |  |  |  |
| Arginine arylamidase | – (0) | NR | NR |
| γ-Glutamyl-transferase | 100 | NR | NR |
| L-Lysine arylamidase | – (0) | NR | NR |
| D-Galactose | – (0) | NR | NR |
| Leucine arylamidase | 100 | +/- | -/v |
| Ellman | 93 | NR | NR |
| Phenylalanine arylamidase | 100 | + | NR |
| L-Proline arylamidase | 100 | + | +/- |
| L-Pyrrolidonyl arylamidase | – (0) | NR | NR |
| Tyrosine arylamidase | 100 | NR | NR |
| Ala-Phe-Pro arylamidase | – (0) | NR | NR |
| D-Glucose | – (0) | – | – |
| Glycogen | – (0) | – | – |
| D-Mannose | – (0) | – | – |
| D-Maltose | – (0) | – | – |
| Saccharose (Sucrose) | – (0) | – | – |
| N-acetyl-D-glucosamine | – (0) | – | – |
| Urease | – (0) | – | + |
| β-galactopyranosidase (Indoxyl) | – (0) | NR | NR |
| Ornithine decarboxylase | 100 | NR | NR |
| α-Glucosidase | – (0) | NR | NR |
| Pyruvate utilisation | 100 | NR | NR |
| Phosphorylcholine | – (0) | NR | NR |
| D-Malate | 100 | +/-v | -/v |
| Maltotriose | – (0) | NR | NR |
| L-Glutamine | 100 | NR | NR |
| Phosphatase | 100 | NR | NR |
| D-Ribose | – (0) | NR | NR |
| D-Xylose | – (0) | – | – |

+, positive; -, negative; v, variable; NR, not reported.

Genomic analysis of *O. otitidis* sp. nov. supported several key phenotypic characteristics including complete absence of urease structural (*ureABC*) and accessory (*ureDEFG*) genes, absence of flagellar biosynthesis genes (including *fli, flg* and *flh* gene clusters), and presence of genes encoding terminal oxidases, including cytochrome oxidase subunits (e.g. *cyoABCD*), and catalase and peroxide detoxification enzymes (e.g. *katA, katE, ahpCF*). *O. otitidis* sp. nov. genomes also encoded multiple components of aerobic respiratory pathways, but lacked complete genes associated with anaerobic respiration, denitrification and enrichment of carbohydrate utilisation pathways - consistent with observed metabolic phenotypes.

### Antibiotic susceptibility

Antibiotics relevant to treatment of otitis media in the Northern Territory, Australia, were chosen for testing. Particular focus was given to ciprofloxacin resistance due to its role as the primary treatment for CSOM in the Northern Territory (as recommended in the otitis media guideline and a local primary healthcare manual)^19^. Antibiotic susceptibility testing of the *O. otitidis* sp. nov. type strain revealed annular radii >6mm for 1.25/23.75μg trimethoprim-sulfamethoxazole disc (10 mm), 10ug gentamicin disc (10 mm), 30ug amoxycillin-clavulanic acid disc (19 mm), 10μg ceftaxidime disc (9 mm) and 10μg polymyxin B disc (8 mm). The strain was resistant to ciprofloxacin (5μg) with no zone of inhibition observed in the disc-based test and an MIC of 32 μg/mL. This resistance profile was consistent across all bar one of the *O. otitidis* sp. nov. isolates (29/30; Table 2). Minimum inhibitory concentrations ranged from <2 to >256 μg/mL (Table 3).

**Table 2:** CDS Disc diffusion results for the 30 *Oligella otitidis* sp. nov. isolates.

| Antibiotic (disc concentration) | MSHR- 50489EDL | All Oo (n=30) |  |
| --- | --- | --- | --- |
|  |  | S | R |
| Ciprofloxacin (5µg) | R | 1 | 29 |
| Trimethoprim-sulfamethoxazole (1.25/23.75µg) | S | 30 | 0 |
| Gentamycin (10µg) | S | 30 | 0 |
| Amoxicillin-clavulanic acid (30µg) | S | 30 | 0 |
| Ceftazidime (10µg) | S | 30 | 0 |
| Polymixin B (10µg) | S | 30 | 0 |

**Table 3:**
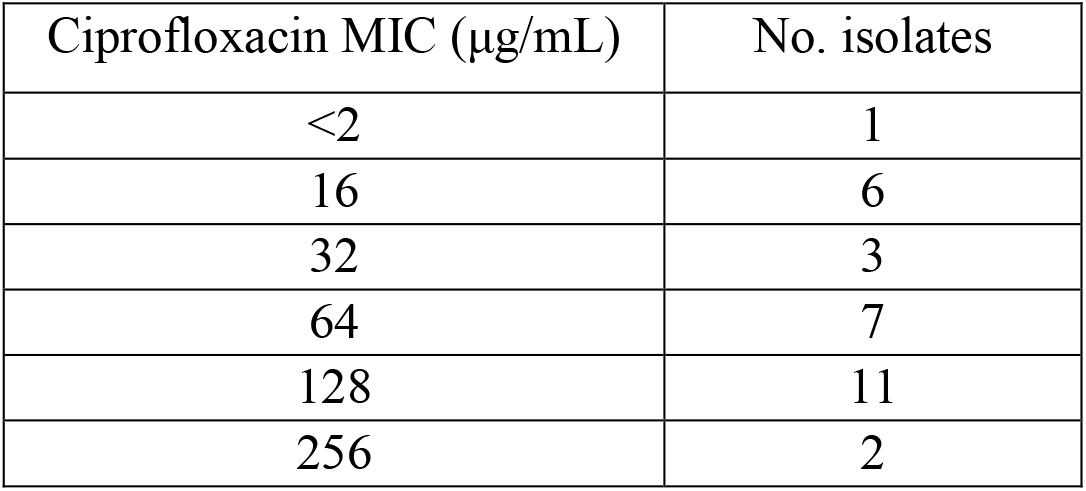
MIC results for the 30 *Oligella otitidis* sp. nov. isolates.

Ciprofloxacin resistance is supported by substitutions in the Quinolone Resistance-Determining Region (QRDR) of *gyrA* (S83I, S83N, S83R, D87N, D87Y) and *parC* (E84Y, E84H, E84V). The presence of multiple mutations at canonical QRDR hotspots is consistent with high-level ciprofloxacin resistance among other members of the Alcaligenaceae family^20,21^. Quinolone resistance (including ciprofloxacin >8->32 μg/mL) is commonly reported among *O. urethralis*-associated infections^22-24^; however, further genome-wide analyses and *in vitro* confirmation is required to fully elucidate quinolone resistance determinants among *Oligella* spp..

### Comparative Genomics

Pangenome analysis of *Oligella* genomes (≥95% amino acid identity) indicated that *O. otitidis* sp. nov. possesses a relatively small core genome of 1,426 genes and a large accessory genome comprising 4,505 genes, consistent with an open pangenome structure. A total of 429 genes were identified that were conserved across *O. otitidis* isolates but absent from *O. urethralis*, while 930 such genes were absent from *O. ureolytica*, indicating substantially greater genomic divergence from the latter species. Of these, 410 genes were not detected in either *O. urethralis* or *O. ureolytica*, representing a distinct accessory genome for *O. otitidis*. Functional categorisation of these genes included conserved hypothetical proteins, with smaller proportions associated with transport and nutrient acquisition, metabolic processes, and cell envelope or surface-associated functions (Table 4), suggesting differences in surface properties and environmental resilience relative to closely related species.

**Table 4.**
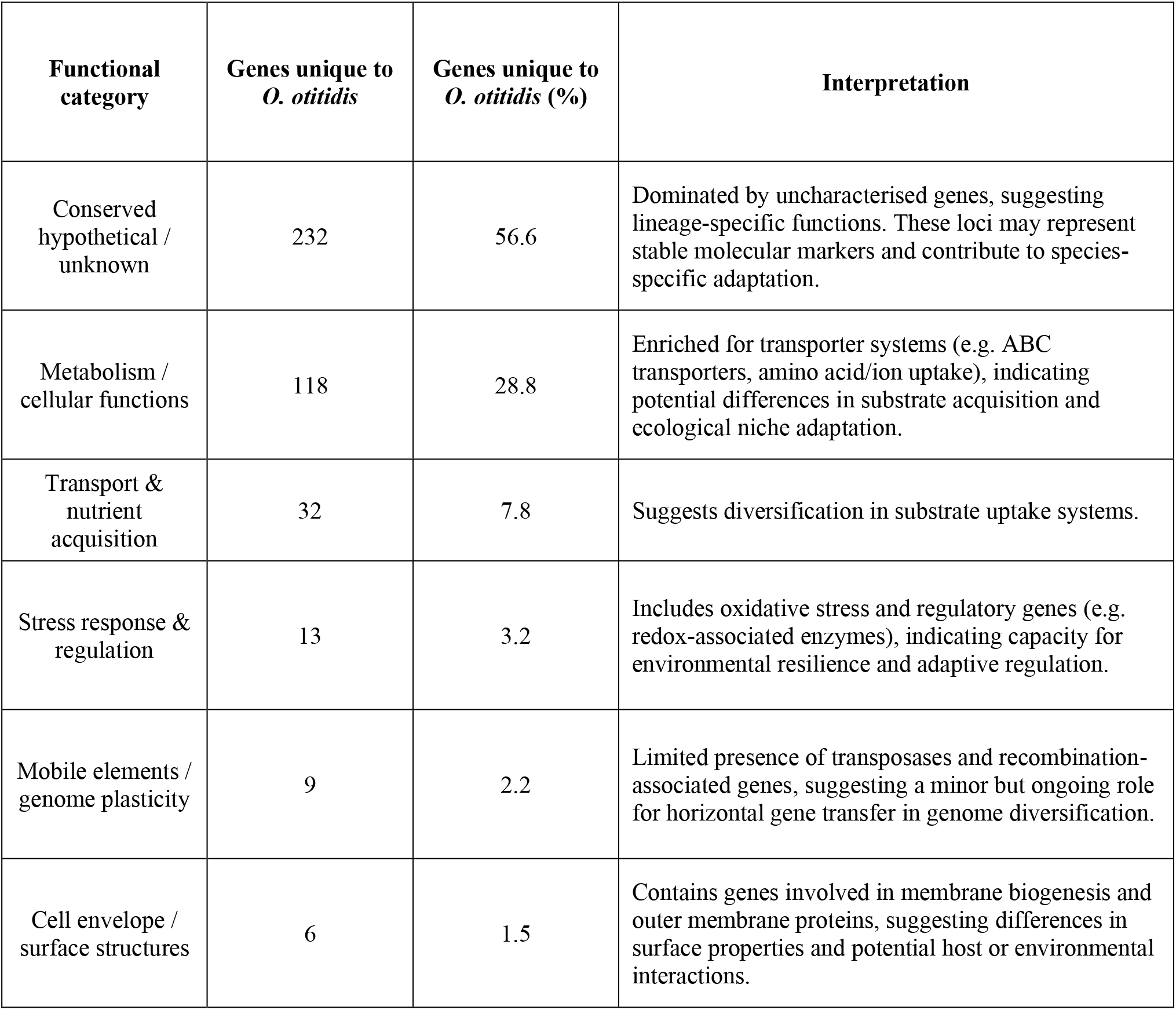
Genes (grouped by function, n= 410) present in *Oligella otitidis* sp. nov. (n=30) and absent in NCBI *O. urethralis* (n=14) and *O. ureolytica* (n=3) genomes, as determined by pangenome analysis. Percentages are calculated relative to the total number of genes uniquely associated with *O. otitidis* sp.nov. (based on ≥95% amino acid identity).

A maximum-likelihood phylogenetic tree based on the core genome alignment (40,528 bp) demonstrated clustering of *Oligella* spp. into distinct species-level lineages (Figure 1). All clinical isolates designate *O. otitidis* sp. nov. formed a single, well-supported monophyletic clade, with high bootstrap support values (≥90–100%) and short branch lengths, indicating high genomic similarity and limited divergence among isolates.

**Figure 1.**
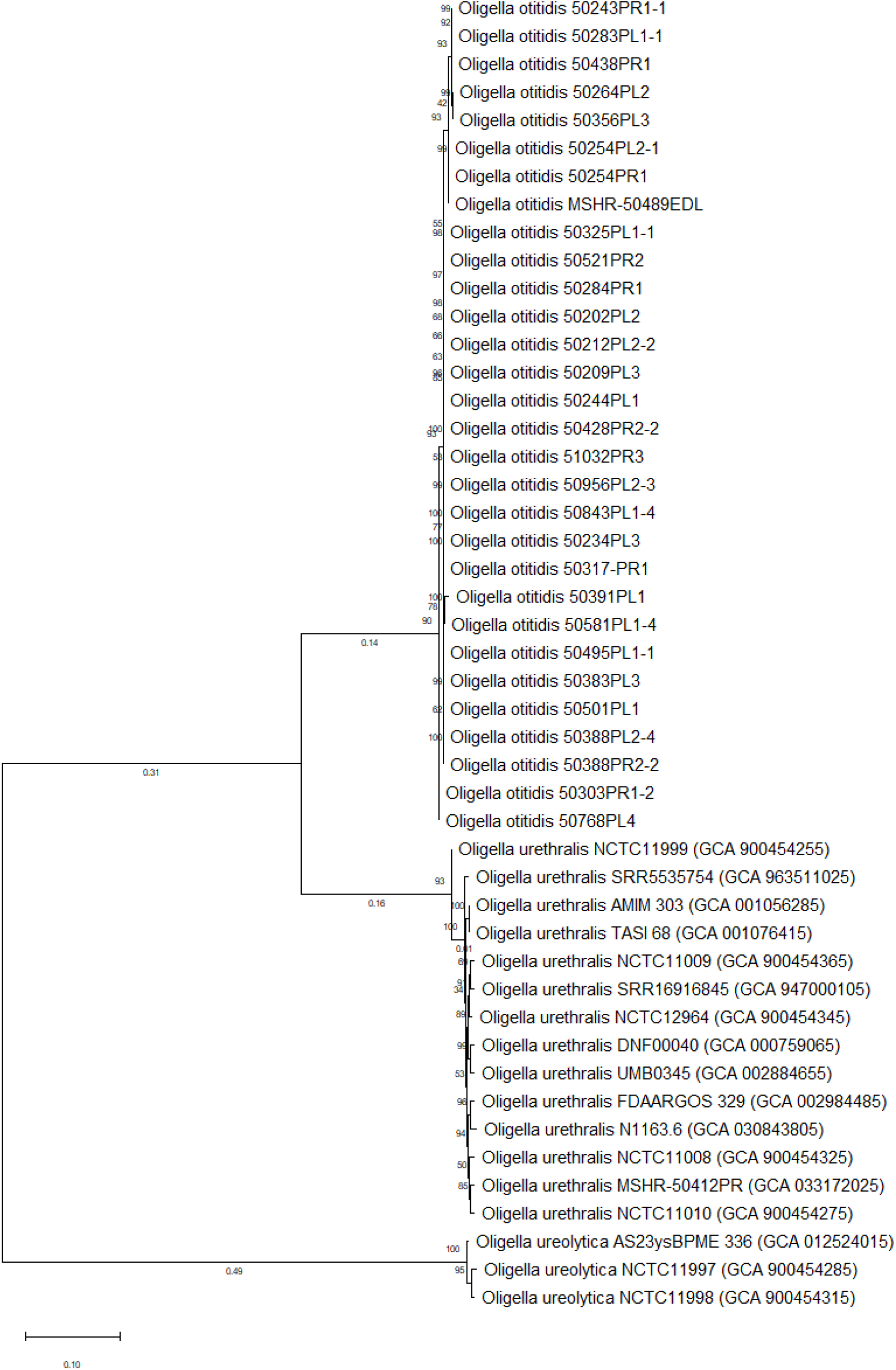
Evolutionary analysis by the Maximum Likelihood method. Maximum-likelihood phylogenetic tree inferred using the General Time Reversible (GTR) model with a discrete Gamma distribution and a proportion of invariant sites (G+I). Bootstrap support values (%) based on 1,000 replicates are shown at branch nodes. The tree is drawn to scale, with branch lengths representing substitutions per site. The analysis included 47 nucleotide sequences.

Average nucleotide identity ([ANI] *O. ureolytica*, 79.70%, alignment coverage 36%; *O. urethralis*, 83%, alignment coverage 63%), digital DNA–DNA hybridization ([dDDh] formula 2, *O. ureolytica*, 24.01%, CI 18.6 – 25.9%; *O. urethralis*, 24.05%, CI 23.4 – 29.5%) and average amino acid identity values (*O. ureolytica*, 80.57 ± 11.25%, orthologous fraction = 65.26%; *O. urethralis*, 84.72 ± 10.67%, orthologous fraction = 72.76) values between the study isolates and *O. ureolytica* and *O. urethralis* are consistent with genus-level relatedness but fall below accepted species thresholds, confirming that *Oligella otitidis* sp. nov. represents a distinct species within the genus (genome statistics shown in Table 5).

**Table 5.** Genome statistics for the 30 *Oligella otitidis* sp. nov. isolates.

| Isolate | Contigs | Size | N50 | GC mol% | CDS | tRNA | rRNA |
| --- | --- | --- | --- | --- | --- | --- | --- |
| 50202 PL-2 | 1 | 2661352 | 2661352 | 45.7 | 2554 | 43 | 9 |
| 50209 PL-3 | 1 | 2679447 | 2679447 | 45.6 | 2506 | 43 | 9 |
| 50212 PL-2(2) | 1 | 2660115 | 2660115 | 45.7 | 2548 | 43 | 9 |
| 50234 PL-3 | 1 | 2614972 | 2614972 | 45.9 | 2382 | 42 | 9 |
| 50243 PR-1(1) | 1 | 2701867 | 2701867 | 45.8 | 2559 | 44 | 9 |
| 50244 PL-1 | 1 | 2729668 | 2729668 | 45.5 | 2577 | 42 | 9 |
| 50254 PL-2(1)(1) | 1 | 2683557 | 2683557 | 45.6 | 2544 | 43 | 9 |
| 50254 PR-1 | 1 | 2688447 | 2688447 | 45.6 | 2615 | 43 | 9 |
| 50264 PL-2 | 1 | 2656863 | 2656863 | 45.8 | 2554 | 43 | 9 |
| 50283 PL-1(1)(1) | 1 | 2724965 | 2724965 | 45.5 | 2641 | 43 | 9 |
| 50284 PR-1 | 1 | 2658457 | 2658457 | 45.7 | 2543 | 43 | 9 |
| 50303 PR-1(2)(1) | 1 | 2695250 | 2695250 | 45.5 | 2586 | 45 | 9 |
| 50317 PR-1 | 1 | 2696810 | 2696810 | 45.5 | 2547 | 43 | 9 |
| 50325 PL-1(1)(1) | 1 | 2593593 | 2593593 | 45.7 | 2438 | 42 | 9 |
| 50356 PL-3 | 1 | 2733800 | 2733800 | 45.6 | 2803 | 42 | 9 |
| 50383 PL-3 | 1 | 2700143 | 2700143 | 45.5 | 2620 | 42 | 9 |
| 50388 PL-2(4) | 1 | 2683850 | 2683850 | 45.9 | 2616 | 42 | 9 |
| 50388 PR-2(2) | 1 | 2668419 | 2668419 | 45.9 | 2551 | 42 | 9 |
| 50391 PL-1 | 1 | 2746971 | 2746971 | 45.5 | 2636 | 44 | 9 |
| 50428 PR-2(2) | 1 | 2697891 | 2697891 | 45.6 | 2575 | 43 | 9 |
| 50438 PR-1 | 1 | 2547072 | 2547072 | 45.9 | 2393 | 43 | 9 |
| 50489 PL-1(3) | 1 | 2733285 | 2733285 | 45.5 | 2598 | 43 | 9 |
| 50495 PL-1(1) | 1 | 2664417 | 2664417 | 45.8 | 2477 | 42 | 9 |
| 50501 PL-1 | 1 | 2687968 | 2687968 | 45.6 | 2587 | 43 | 9 |
| 50521 PR-2 | 1 | 2762983 | 2762983 | 45.5 | 2628 | 44 | 9 |
| 50581 PL-1(4) | 1 | 2664144 | 2664144 | 45.6 | 2487 | 43 | 9 |
| 50768 PL-4 | 1 | 2679466 | 2679466 | 45.5 | 2813 | 45 | 9 |
| 50843 PL-1(4) | 1 | 2689945 | 2689945 | 45.7 | 2533 | 44 | 9 |
| 50956 PL-2(3) | 1 | 2668209 | 2668209 | 45.7 | 2508 | 43 | 9 |
| 51032 PR-3 | 1 | 2636435 | 2636435 | 45.7 | 2494 | 44 | 9 |

We consequently classify this strain as a new species of the phylum *Pseudomonadota*, Class *Betaproteobacteria*, Order *Burkholderiales*, family *Alcaligenaceae* and Genus *Oligella*, and propose the name *Oligella otitidis* sp. nov. (o.ti’ti.dis. N.L. gen. n. *otitidis*, of otitis, of inflammation of the ear). The complete average genome is 2.67 Mbp (2,565 coding sequences, 45.69 % GC content, 43 tRNA and 9 rRNA).

### *Oligella otitidis* sp. nov. species description

*Oligella otitidis* sp. nov. is an aerobic Gram-negative, oxidase- and catalase-positive, non-spore-forming, non-motile cocco-bacillus. The optimum range for growth is pH 8–9, NaCl concentrations 4–5%, 25–42°C. *O. otitidis* produces small, circular, smooth, whitish-opaque and occasionally mucoid colonies after 48-hour incubation on horse blood or MacConkey agar in aerobic conditions. Enzymatic activities are positive for γ-Glutamyl-transferase, L-Proline arylamidase, leucine arylamidase, tyrosine arylamidase, ornithine decarboxylase and phenylalanine arylamidase and negative for arginine arylamidase, L-Lysine arylamidase, L-Pyrrolidonyl arylamidase, urease, α-Glucosidase and Ala-Phe-Pro arylamidase. Carbohydrate fermentation is negative for glucose, glycogen, mannose, maltose, sucrose, N-acetyl-D-glucosamine and D-Galactose. *O. otitidis* MSHR-50489EDL (ATCC: TSD-462; DSMZ: DSM 118617), has an ANI <83.1% similar to the phylogenetically closest related species, *O. urethralis* and *O. urealytica*, and is proposed as the type strain of this new species. The type strain and all isolates recovered to date are from ear discharge of children with CSOM. The complete average genome is 2.67 Mbp (2,565 coding sequences, 45.69 % GC content, 43 tRNA and 9 rRNA).

### Nucleotide sequence accession number

The *O. otitidis* type strain genome sequence was deposited in Genbank under accession number CP158480.

### Deposit in culture collections

Strain MSHR-50489EDL has been deposited in the ATCC (ATCC: TSD-462) and DSMZ (DSM 118617) isolate collections.

## Acknowledgements

We would like to acknowledge Christine Wigger, Deborah Taylor-Thompson, Katrina Lawrence and the families who participated in the IHEARBETA randomized controlled trial.

The study was approved by Human Research Ethics Committee of Northern Territory Health and Menzies School of Health Research (Approval #2014-2170).

The study was conceptualised by JB and RLM who wrote the original manuscript with BA. Funding was acquired by JB, RLM, AJL, PSM, HSV and BA. Laboratory methods were designed by JB, RLM, PKM, BA. Laboratory investigations were undertaken by JB, AC, PKM, BA, NG, BJ, VG and MB, and validations by VR, BA, BH, NG, MB. Isolation of Oligella strains was done by PKM and AC under the supervision of JB and RLM. The project and resources were administered by JB, BA and PKM. Student supervision was undertaken by JB, RLM, PKM and BA. Software responsibilities were JB, PKM, RLM, and BA. Data Curation was undertaken by JB, AC, PKM, BA, NG, BH and MB. Type strain NCBI submission was done by PKM. Majority of analysis and visualisation was done by BA. All authors reviewed the manuscript and approved submission.

## Reference list

1. Freese HM, Meier-Kolthoff JP,Sardà Carbasse J, Afolayan Ayorinde O, Göker M. TYGS and LPSN in 2025: a Global Core Biodata Resource for genome-based classification and nomenclature of prokaryotes within DSMZ Digital Diversity. Nucleic Acids Research 2026; 54(D1): D884–D91.

2. Serandour P, Plouzeau C, Michaud A, et al. The First Lethal Infection by Oligella ureolytica: A Case Report and Review of the Literature. Antibiotics (Basel) 2023; 12(9).

3. Farfour E, Vasse M, Vallée A. Oligella spp.: A systematic review on an uncommon urinary pathogen. Eur J Clin Microbiol Infect Dis 2024; 43(6): 1037–50.

4. Santos-Cortez RL, Hutchinson DS, Ajami NJ, et al. Middle ear microbiome differences in indigenous Filipinos with chronic otitis media due to a duplication in the A2ML1 gene. Infectious diseases of poverty 2016; 5(1): 97.

5. Taylor SL, Papanicolas LE, Richards A, et al. Ear microbiota and middle ear disease: a longitudinal pilot study of Aboriginal children in a remote south Australian setting. BMC Microbiol 2022; 22(1): 24.

6. Mandal PK, Cleanthous A, Rigas V, et al. Complete genome sequence of Oligella urethralis MSHR-50412PR, isolated from an ear discharge swab of a child with chronic suppurative otitis media. Microbiol Resour Announc 2024; 13(2): e0107123.

7. Wigger C, Leach AJ, Beissbarth J, et al. Povidone-iodine ear wash and oral cotrimoxazole for chronic suppurative otitis media in Australian aboriginal children: study protocol for factorial design randomised controlled trial. BMC pharmacology & toxicology 2019; 20(1): 46.

8. QIAamp® DNA Mini and Blood Mini Handbook 5ed: QIAGEN; 2016.

9. Bolger AM, Lohse M, Usadel B. Trimmomatic: a flexible trimmer for Illumina sequence data.Bioinformatics 2014; 30(15): 2114–20.

10. Wick RR, Judd LM, Gorrie CL, Holt KE. Unicycler: Resolving bacterial genome assemblies from short and long sequencing reads. PLoS Comput Biol 2017; 13(6): e1005595.

11. Dong MJ, Luo H, Gao F. Ori-Finder 2022: A Comprehensive Web Server for Prediction and Analysis of Bacterial Replication Origins. Genomics Proteomics Bioinformatics 2022; 20(6): 1207–13.

12. Carver T, Harris SR, Berriman M, Parkhill J, McQuillan JA. Artemis: an integrated platform for visualization and analysis of high-throughput sequence-based experimental data. Bioinformatics 2012; 28(4): 464–9.

13. Meier-Kolthoff JP, Auch AF, Klenk H-P, Göker M. Genome sequence-based species delimitation with confidence intervals and improved distance functions. BMC Bioinformatics 2013; 14(1): 60.

14. Meier-Kolthoff JP, Carbasse JS, Peinado-Olarte RL, Göker M. TYGS and LPSN: a database tandem for fast and reliable genome-based classification and nomenclature of prokaryotes. Nucleic Acids Research 2021; 50(D1): D801–D7.

15. Nei M, Kumar S. Molecular Evolution and Phylogenetics: Oxford University Press; 2000.

16. Felsenstein J. CONFIDENCE LIMITS ON PHYLOGENIES: AN APPROACH USING THE BOOTSTRAP.Evolution 1985; 39(4): 783–91.

17. Saitou N, Nei M. The neighbor-joining method: a new method for reconstructing phylogenetic trees. Molecular Biology and Evolution 1987; 4(4): 406–25.

18. Kumar S, Stecher G, Suleski M, Sanderford M, Sharma S, Tamura K. MEGA12: Molecular Evolutionary Genetic Analysis Version 12 for Adaptive and Green Computing. Molecular Biology and Evolution 2024; 41(12).

19. Leach AJ, Morris PS, Coates HL, et al. Otitis media guidelines for Australian Aboriginal and Torres Strait Islander children: summary of recommendations. Med J Aust 2021; 214(5): 228–33.

20. Furlan JPR, Sanchez DG, Gallo IFL, Stehling EG. Replicon typing of plasmids in environmental Achromobacter sp. producing quinolone-resistant determinants. APMIS : acta pathologica, microbiologica, et immunologica Scandinavica 2018; 126(11): 864–9.

21. Magallon A, Roussel M, Neuwirth C, et al. Fluoroquinolone resistance in Achromobacter spp.: substitutions in QRDRs of GyrA, GyrB, ParC and ParE and implication of the RND efflux system AxyEF-OprN. J Antimicrob Chemoth 2020; 76(2): 297–304.

22. Muranaka E, Kurihara M, Okawa N, et al. Oligella urethralis bacteremia associated with urinary tract obstruction: A case report and literature review. Journal of Infection and Chemotherapy 2026; 32(2): 102909.

23. Beauruelle C, Le Bars H, Bocher S, Tande D, Hery-Arnaud G. The Brief Case: Extragenitourinary Location of Oligella urethralis. J Clin Microbiol 2019; 57(8).

24. Riley UB, Bignardi G, Goldberg L, Johnson AP, Holmes B. Quinolone resistance in Oligella urethralis-associated chronic ambulatory peritoneal dialysis peritonitis. J Infect 1996; 32(2): 155–6.

